# DINOcell: learning generalizable perturbation effects through self-distillation

**DOI:** 10.64898/2025.12.16.694747

**Authors:** Jonathan Li, Nilgun Tasdemir, Sofia Kyriazopoulou Panagiotopoulou, Sophie Xu, Allyson Merrell, Josephine Susanto, Ashley Cass, David DeTomaso, Holly Steach, Julie Chow, Li Wang, Neal Ravindra, Aram Kang, Grace Zheng, Mary Chua, Per Knudsgaard, Emmie He, Fabio Ingrao, Angela Boroughs, Brie Copperman, Nia Charrington, Kathryn Abboud, Lionel Berthoin, Daniel Baca, Lina Leon, Jessica Kotov, Glenn Wozniak, Andrew Cardozo, Kristina Vucci, Adam Barner, Charles Yi, Catherine Oh, Mandi Simon, Saurabh Paliwal, Matt Drever, Brendan Galvin, Michelle Tan, Gustavo Guzman, Kevin Loh, Azalea Ong, Marian Sandoval, Meng Lim, Emily Ng, Brishette Lincoln-Cabatu, Jimmy Wu, Ankim Nguyen, Mehal Patel, Andrea Fua, Kevin Wong, Jon Chen, Ed Yashin, Terrence Yim, Aaron Cooper, Luke Cassereau, W. Nicholas Haining, Amaro Taylor-Weiner

## Abstract

Predicting cellular responses to therapeutics is a promising approach for novel target discovery. However, state-of-the-art computational models designed to predict perturbation effects struggle to generalize and outperform simple baselines. We present DINOcell, a weakly supervised framework that adapts self-distillation to single-cell transcriptomics for predicting perturbation effects. We demonstrate that DINOcell outperforms baselines in predicting the effects of single gene perturbations. Furthermore, DINOcell accurately predicts non-additive effects of combination perturbations, indicating its capacity to model complex genetic interactions. Finally, we show that DINOcell learns representations that capture biological signals and is a promising, generalizable approach for *in silico* perturbation modeling, providing a valuable tool for accelerating therapeutic target discovery.

## 1 Introduction

A fundamental challenge in therapeutic development is predicting how complex biological systems will respond to interventions or perturbations. The ability to infer the effects of untested perturbations could greatly accelerate the discovery of drug targets that produce therapeutically desirable cellular phenotypes. This is especially valuable when experimentally testing all possible genetic or molecular interventions (and their combinations) is impossible due to physical limitations. These experimental limitations create the need for computational models that can generalize from available data to predict the effects of unseen perturbations *in silico*, enabling search beyond physical constraints. However, current deep learning models struggle to disentangle the effects of perturbations on cell state from technical variation and thus underperform simple baselines [1, 2].

Single-cell RNA sequencing (scRNA-seq) data has a spectrum of variation by nature. The largest sources of variation can be biological (e.g. cell type, donor) or technical (e.g. sequencer, experimental conditions). In contrast, perturbations (e.g. CRISPR knockouts), even those that have significant biological impact, often have narrower transcriptional effects with lower variance that can affect a limited number of genes. Learning generalizable representations that can encode variation across scales, particularly small sources of variation induced by perturbations, is a key challenge. Capturing variation on this scale is essential because it allows us to predict how cells within a particular state will respond to targeted perturbations, which is a more advanced goal than, say, learning to differentiate between major cell groups (e.g., neurons vs. immune cells).

Self-supervised learning can learn semantically meaningful representations that are resilient to technical variations without the need for supervised labels. DINO (Distillation with No Labels), a self-supervised model first introduced for computer vision, leverages the concept of invariances to learn representations that generalize across diverse downstream tasks including classification, segmentation, and open-set object detection [3]. DINO uses a self-distillation framework, training a student model to match the teacher’s output from different augmented views of the same input. This process encourages learned features that are invariant to noise, thereby encoding content-specific information. DINO employs centering and sharpening to structure the latent space and avoid learning a trivial solution in which all inputs collapse to a single representation (mode collapse). Sharpening applies a lower softmax temperature to the teacher’s outputs, encouraging the student to learn more discriminative and diverse representations. Notably, unlike contrastive approaches, DINO does not require negative class labels. Considering the prevalence of technical variation (e.g. batch effects) and the scarcity of high-quality labels in single-cell data, we propose that a similar approach holds promise for the single-cell domain.

We introduce DINOcell, an adaptation of DINO, for predicting the effects of CRISPR perturbations on single-cell transcriptomes. We train the model through two tasks: (1) perturbation of a control cell; and (2) alignment of cells that receive the same perturbation. The rest of the manuscript is organized as follows: First, we provide an overview of the architecture and training objectives (Section 2.1). Then, we compare the model against baseline methods for predicting the effects of single and combination CRISPR guides in immortalized human cell lines and primary human T cells (Sections 2.2 and 2.3). We show that DINOcell can predict the effects of unseen non-additive combinations (e.g. in genes with epistatic interactions) (Section 2.3). Finally, we demonstrate that embeddings generated by DINOcell are semantically meaningful and have improved generalizability between cellular contexts (Section 2.4).

## 2 Results

### 2.1 DINOcell for predicting perturbation effects via self-distillation

DINOcell extends the self-distillation framework of DINO [3] to scRNA-seq data. In DINO, a student network is trained to match the teacher’s outputs, where the teacher is an exponential moving average (EMA) of the student (Figure 1A). This enables self-supervised representation learning from unlabeled data by forcing the student to predict the teacher’s sharpened softmax outputs (predicted pseudolabels) across multiple augmented views of the same input. Two mechanisms are incorporated to prevent mode collapse. Centering subtracts a running mean of the teacher’s logits, removing distributional drift and discouraging dominance of any single dimension. Sharpening applies a lower softmax temperature to the teacher’s outputs, encouraging the student to learn more discriminative and diverse representations.

**Figure 1:**
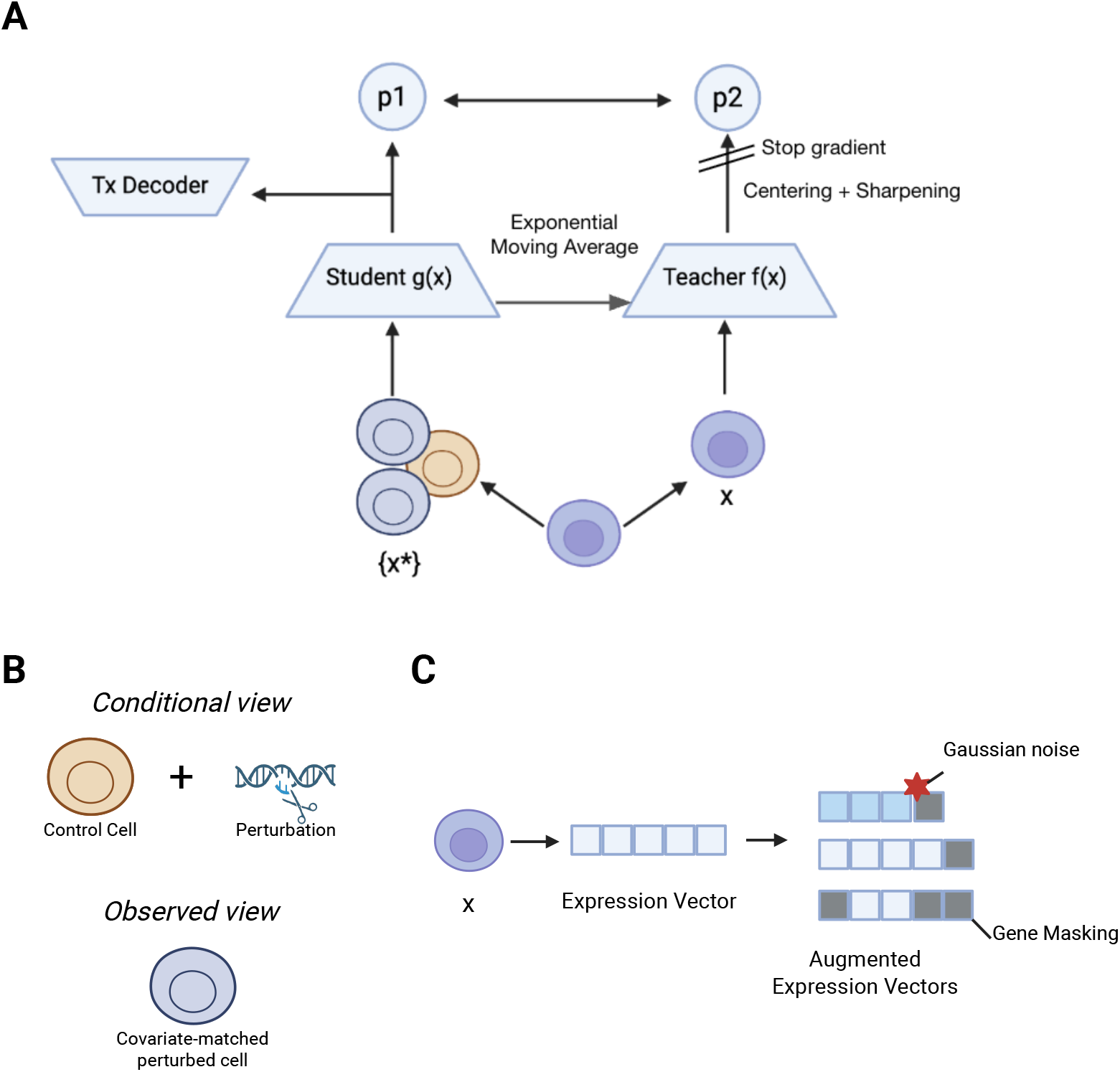
Method & Dataset Overview. A) DINOcell schematic: training of a student model through self-distillation, utilizing a DINO objective calculated from predicted pseudolabels (p1 and p2). A teacher model encodes a vector pair of gene expression and gene ID tokens from a single cell (*x*). The student model then encodes augmented views ({*x*\*}) of *x*, with its aligned representation acting as an information bottleneck. This bottleneck is subsequently used by the Tx Decoder to predict the expression of perturbed cells. All models are parameterized as transformer models. B) Student view types schematic: types of inputs provided to the student model. Given a perturbed cell *x*, we construct several views, {*x*\*}. These are configured to be drawn from two types: conditional and observed views. In the conditional view, the student is given a learnable gene perturbation representation with expression from a control cell. In the observed view, we select a different cell from the same donor and cell type as *x* that received the same CRISPRi guide. Note the teacher only receives unaugmented cells (*x*). C) Student view augmentations: We add augmentations to student views. We leverage two types of augmentations: addition of Gaussian noise to expression values and aggressive random masking of genes.

We extend DINO to cells by operating on single-cell transcriptomes. Each cell (*x*) is tokenized at the gene level, such that *x* = {*e*_*i*_ }, where *e*_*i*_ is the embedding for gene *i*. The embedding is an element-wise sum of an expression value embedding and a learnable gene name embedding, similar to scGPT [4]. A CLS token is appended to *x* to capture a global cell representation. Following DINO, we generate multiple augmented views {*x*\*} of each cell by downsampling gene tokens and adding Gaussian noise to expression values (Figure 1B,C). The teacher processes an unaugmented cell, while the student processes augmented ones (described below). The student’s predicted pseudolabel distributions are trained to match that of the teacher using cross-entropy loss across all student-teacher pairs. Here, pseudolabels can be loosely interpreted as a soft-classification of emergent clusters of cells. The student is updated via gradient descent, and the teacher’s weights are updated by EMA, ensuring a stable and evolving target throughout training. The student encoder is initialized from a transformer-based pretrained encoder-decoder model (88 million parameters) with a masked token objective and trained on 75.1M single-cell transcriptomes drawn from both public and internal datasets.

The self-distillation loss encourages representation learning. To obtain expression values from cell representations, we include an auxiliary expression-decoder head attached to the student CLS token. The decoder uses FiLM conditioning [5], where the student CLS token generates feature-wise affine parameters (scale and shift) to modulate the gene token embeddings via an element-wise product. The modulated embeddings are then projected (via a dense projection) to predict expression values. The resulting mean squared error (MSE) loss is scaled and added to the DINO objective

To model perturbation effects, DINOcell introduces two complementary student inputs that jointly capture both conditional and observed mappings of perturbation outcomes. In the conditional view, the student receives: (i) the tokenized expression of a control cell^1^; and (ii) one or more perturbation tokens encoding the targeted gene(s) (Figure 1B). Each perturbation token is produced by an adapter layer that projects the GenePT embedding [6] of the targeted gene through a Gaussian error linear unit (GELU; [7])-activated multilayer perceptron to align it with the student encoder’s dimensionality. The adapted perturbation token(s) are prepended to the control cell’s tokens before being passed to the student encoder. In the observed view, the student instead receives the augmented expression profile of a covariate-matched perturbed cell directly. Including the observed view provides a stabilizing and anchoring signal, ensuring that representations inferred from conditional views remain consistent with those learned directly from measured perturbed expression. Both configurations are trained against the same teacher target, with both contributing equally to the DINO loss during optimization.

Finally, to reduce covariate bias in heterogeneous single-cell datasets, DINOcell replaces DINO’s single global center with conditional centering. A single global center tends to bias representations toward dominant covariates (e.g. donor, cell type, or experiment), causing a form of mode collapse in which representations primarily organize along these axes. By maintaining a center for each covariate group, conditional centering allows the model to focus on finer perturbation-driven variation. Empirically, conditional centering yields more stable training and improved expression-level prediction accuracy (data not shown).

### 2.2 DINOcell accurately predicts perturbation effects across cell contexts

We evaluated DINOcell on a generalization task designed to assess its ability to predict perturbation effects in partially unseen cell contexts (e.g. new experimental system, different cell type, etc.). Specifically, we trained DINOcell with 30% of perturbations in a target context and asked how well the models could predict the remaining 70% of perturbations (Figure 2A). The primary metric we used for evaluation in this work is the delta Pearson correlation coefficient (dPCC), computed as the Pearson correlation between true and predicted pseudobulked fold-changes (logFC), both relative to the true unperturbed control.

**Figure 2:**
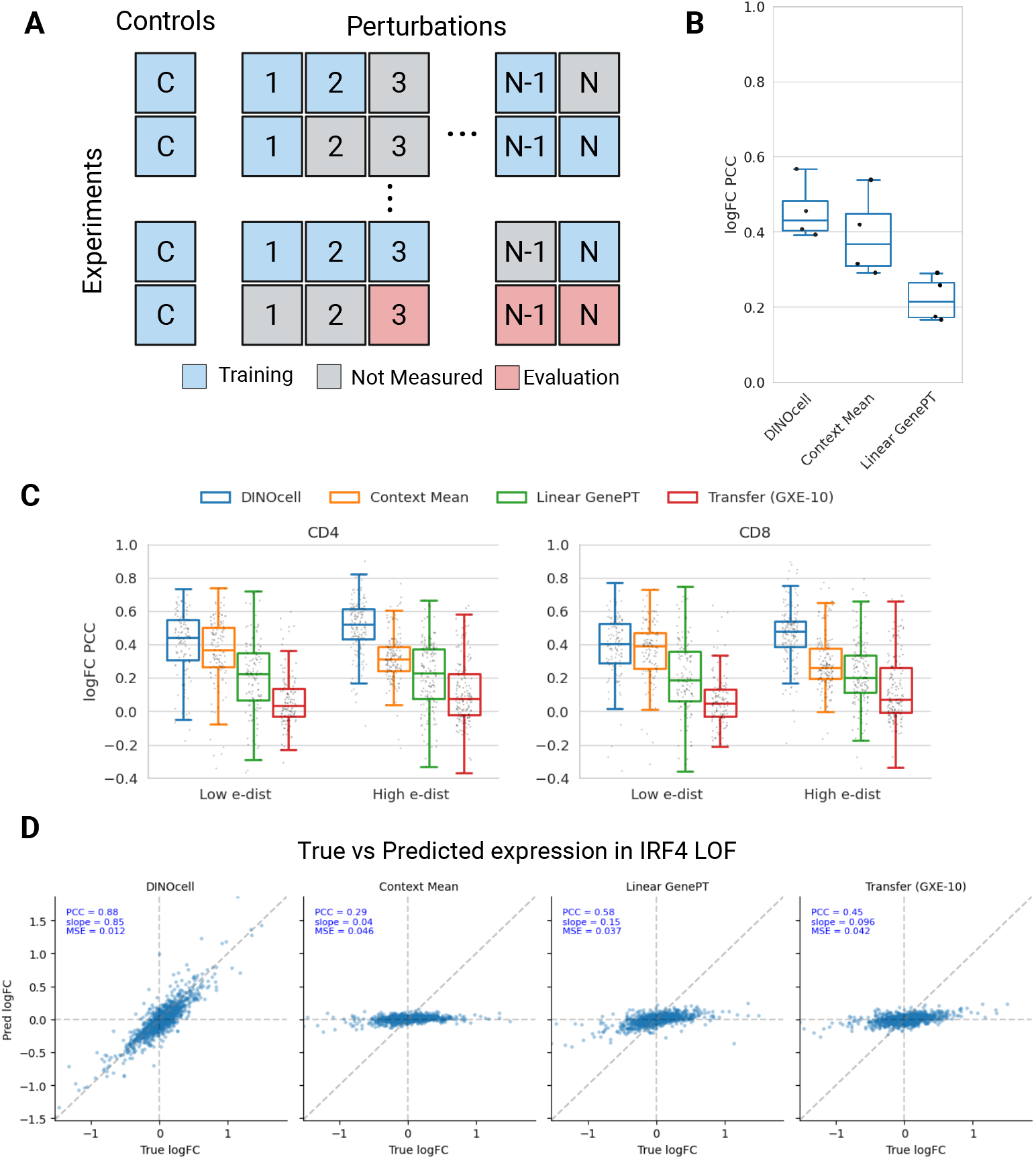
Results Summary for Single Perturbation Effect Prediction. A) Schematic of context transfer task for single perturbations: A context refers to a cell line and/or experiment. In each run, 70% of the available perturbations are held out as the test set from a single target context. All perturbations from the other contexts, as well as the remaining 30% of perturbations from the target context, are used for training. B) Context transfer results for Replogle-Nadig dataset: Boxplots (boxes = IQR, whiskers = 95% CI, lines = median) summarizing model performance (logFC PCC) across data splits. Each point represents a split in which one of the 4 cell lines in Replogle-Nadig was chosen as the target context. C) Context transfer results for internal datasets: gxe63 was chosen as the target context. Each point represents the logFC PCC of an individual perturbation in CD4 or CD8 cells. Perturbations with low and high e-distances (log1p(e-dist) > 2) are plotted separately. D) Comparison of predictions for *IRF4*-targeted CRISPRi: Each panel shows the predicted vs. true logFC expression for a different model or baseline. Each point is a single gene.

We benchmarked DINOcell’s performance against several baselines. First, the context mean baseline predicts each perturbation’s expression by averaging all training perturbations within that context. Second, the linear GenePT baseline uses pretrained GenePT embeddings as input features and learns a context-specific linear mapping to gene expression which is then applied to unseen perturbations. All models, including DINOcell, were trained to predict log-normalized expression for the top 2,000 highly variable genes (HVGs).

We first applied this approach to the Replogle-Nadig genetic perturbation dataset [8, 9] across four cell lines, filtering out perturbations with low on-target efficacy (2,024 remaining). We ran four iterations in which one cell line served as the target, with 70% of its perturbations withheld for evaluation. DINOcell achieved significantly higher mean dPCC scores than both baselines for each target cell line (Figure 2B, DINOcell vs. context mean p-val = 0.055; DINOcell vs. linear GenePT p-val = 0.0011).

We next tested generalization on an internal dataset in primary human T cells that were polarized *in vitro* to adopt a Th17/Tc17 phenotype (see Methods). This Th17/Tc17 perturbation dataset was composed of a genome-wide loss-of-function (LOF) perturb-seq screen (gxe10; 18,690 perturbations) and two curated sub-genome-scale LOF screens (gxe46, 178 perturbations; and gxe63-singles, 736 perturbations) (Supplementary Figure 1A,B). We followed a similar split procedure as before, holding out 70% of LOF singles in gxe63 and training on the rest of the data, excluding any combination guide cells. For this analysis we included a third baseline, namely a log fold-change (logFC) transfer baseline in which the logFC from gxe10 is applied to gxe63, matching by cell type (i.e. CD4 or CD8 T cell).

We found that DINOcell again outperformed all three baselines in both CD4 and CD8 T cells, with predicted logFC profiles showing higher correlation to ground truth (Figure 2C). DINOcell outperformed baselines regardless of the magnitude of effect size of the perturbations, as measured by the e-distance metric [8]. To more closely study the predictive performance of DINOcell we next compared predictions for the effects of *IRF4* LOF on Th17 T cells. *IRF4* is crucial for fate determination of Th17 cells [10], and is one of the highest effect perturbation targets in our dataset. DINOcell captures the effect of this perturbation much more accurately than baseline methods (Figure 2D; DINOcell vs. Linear baseline PCC: 0.88 vs. 0.58). These results demonstrate that DINOcell effectively generalizes single-target perturbations to cellular contexts with sparse coverage.

### 2.3 DINOcell can predict non-additive effects of perturbation combinations

Next, we evaluated DINOcell on predicting the effects of perturbation combinations in Th17 cells, where two LOF perturbations were introduced into each cell. We generated a panel of two-target perturbations using 33 single-gene knockouts from those targets used in the gxe63-singles library, resulting in 528 possible combinations (gxe63-combos). Training data included gxe10, gxe46, and gxe63-singles, plus 26 combinations (5% of total) from gxe63-combos to expose the model to examples of combinatorial effects. The remaining 95% of combinations were held out for evaluation (Figure 3A).

**Figure 3:**
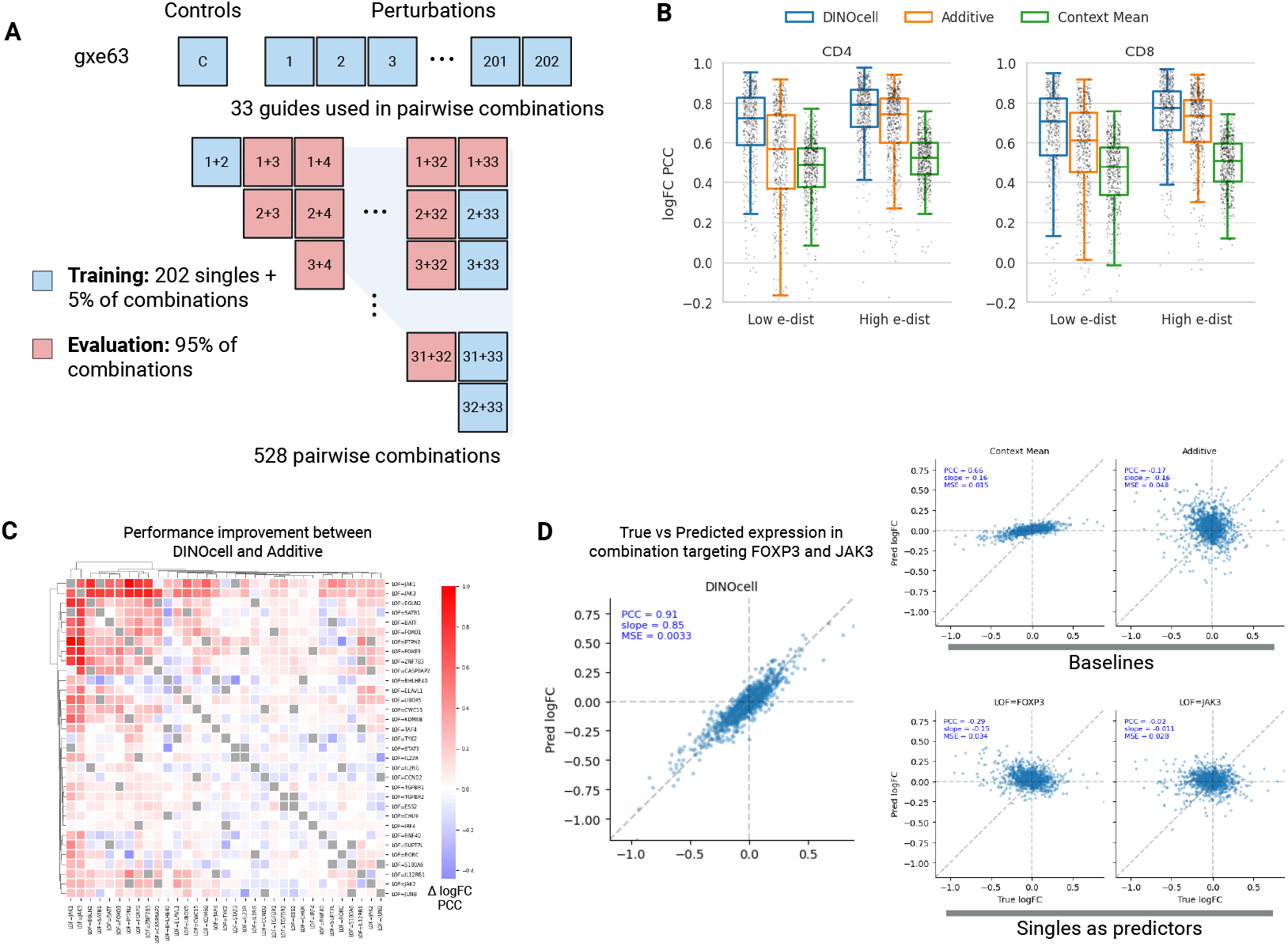
Results Summary for Combination Perturbation Effect Prediction. A) Overview of data splits for evaluation on unseen combinations: gxe63 is composed of 202 single and 528 combination perturbations performed in 4 donors. Pairwise combinations were generated from 33 selected singles (*C*(33, 2) = 528). All singles and 5% of the combinations were used for training. The remaining 95% of combinations were used for evaluation. B) Results for prediction of unseen paired perturbation effects: Boxplots (boxes = IQR, whiskers = 95% CI, lines = median) summarizing model performance (logFC PCC) for the held-out combinations. Each point represents the logFC PCC of an individual perturbation in CD4 or CD8 cells. Perturbations with low and high e-distances (log1p(e-dist) > 2) are plotted separately. C) Improvement in performance between DINOcell and the additive baseline: Each square shows the difference in logFC PCC between DINOcell and the additive baseline for individual combinations. Red indicates higher DINOcell performance, blue indicates higher additive performance; combinations seen during training are greyed out. D) Predicted vs. true logFC expression for an example non-additive combination (LOF=*FOXP3*|*JAK3*): The top row shows DINOcell and baseline predictions; the bottom row shows the individual single-gene perturbations (LOF=*FOXP3*, LOF=*JAK3*). Each point is a single gene.

We compared the performance of DINOcell to that of an additive baseline, in which the predicted logFC for a combination was computed as the sum of the pseudobulked fold changes for its individual components (e.g., *f* (*X, Y*) = *f* (*X*) + *f* (*Y*)). DINOcell predictions were substantially more accurate in both CD4 and CD8 cells, outperforming the additive baseline for 75% of combinations in CD4 and 67% in CD8 (Figure 3B). Again, these performance improvements were seen in both weak- and strong-effect perturbations.

Finally, we evaluated whether DINOcell could accurately predict the effects of perturbation combinations that showed non-additive effects. Capturing these non-additive, or epistatic, interactions is a major challenge for cell models, since they reflect complex regulatory dependencies that cannot easily be inferred from single-gene perturbations alone. In our experimental dataset, we noticed that combinations targeting certain genes (*JAK1, JAK3, FOXP3*, and *PTPN2*), which interact through the JAK-STAT signaling pathway, produced non-additive effects. Despite having no explicit training to indicate which interactions were non-additive, DINOcell was able to accurately predict the effects of these combinations (Figure 3C). For instance, the *FOXP3*|*JAK3* combination exhibits fold-change patterns uncorrelated with either single perturbation and poorly predicted by the additive model (dPCC=0.13). The context-mean baseline shows moderate correlation (dPCC=0.66) but severely underestimates effect magnitude (slope=0.16). In contrast, DINOcell achieved high accuracy (dPCC=0.91; Figure 3D).

To rule out the possibility that the model had simply memorized the effects of another combination seen during training, we clustered perturbations to identify those with similarity to *FOXP3*|*JAK3* (Supplementary Figure 2A). We found 4 combinations (LOF=*JAK1*|*STAT3*, LOF=*JAK1*|*SATB1*, LOF=*EGLN2*|*IL2RG*, LOF=*IL2RG*|*RORC*) that were seen during training and whose pseudobulked expression cluster with LOF=*FOXP3*|*JAK3* in either CD4 or CD8 cells (Supplementary Figure 2B). While these combinations were highly correlated with LOF=*FOXP3*|*JAK3* (dPCC=0.69– 0.92), the individual components of these combinations were generally poorly correlated with LOF=*FOXP3* and LOF=*JAK3*, suggesting that the model is not performing a trivial mapping (Supplementary Figure 2C). For example, LOF=*JAK1* (part of LOF=*JAK1*|*STAT3*) was highly correlated with LOF=*JAK3* (CD4 dPCC=0.78, CD8 dPCC=0.85). If the model had learned a trivial mapping, we would expect LOF=*FOXP3* to be correlated with LOF=*STAT3*, yet we find that they are poorly correlated (CD4 dPCC=0.022, CD8 dPCC=0.27).

Together, these results indicate that DINOcell captures genuine non-additive combinatorial effects, suggesting it learns biologically meaningful representations of complex cellular responses to perturbation rather than relying on simple, linear correlations.

### 2.4 DINOcell produces high-quality embeddings

We wanted to assess whether DINOcell was able to generate high-quality single-cell embeddings, as a measure of its ability to learn biologically meaningful representations. First, we verified whether the model was able to align perturbed representations from both the “observed view” (expression from a covariate-matched cell) as well as the “conditional view” (control expression + perturbation label). For both views, we sampled 20 cells per perturbation within each covariate group and took the average of their embeddings. Plotting them together on a UMAP shows that the observed views cluster closely with the corresponding conditional views (Supplementary Figure 3A,B). This indicates that the model successfully learns a latent space shared by both views, enabling the use of a common expression decoder.

Next, we investigated the extent to which known sources of variation, like donor, cell type and perturbation, are encoded in the embeddings. We reasoned that the observed views should encourage the model to learn invariances within a covariate group and produce more separable embeddings relative to the pretrained model. To test this, we prompted the student model with unaugmented perturbed cells and extracted their CLS token embeddings (Figure 4A). For cells sampled from gxe63, we find that DINOcell produces more separable embeddings compared to pretrained and raw expression embeddings (Figure 4B). We quantified this separability by measuring the frequency that the closest neighbor for each cell came from the same covariate group or perturbation (k-NN top-1 purity), normalized to a shuffled baseline (n=100). We found that indeed, DINOcell embeddings were more coherent with respect to covariates and perturbation labels compared with the pretrained model and raw expression (Figure 4C).

**Figure 4:**
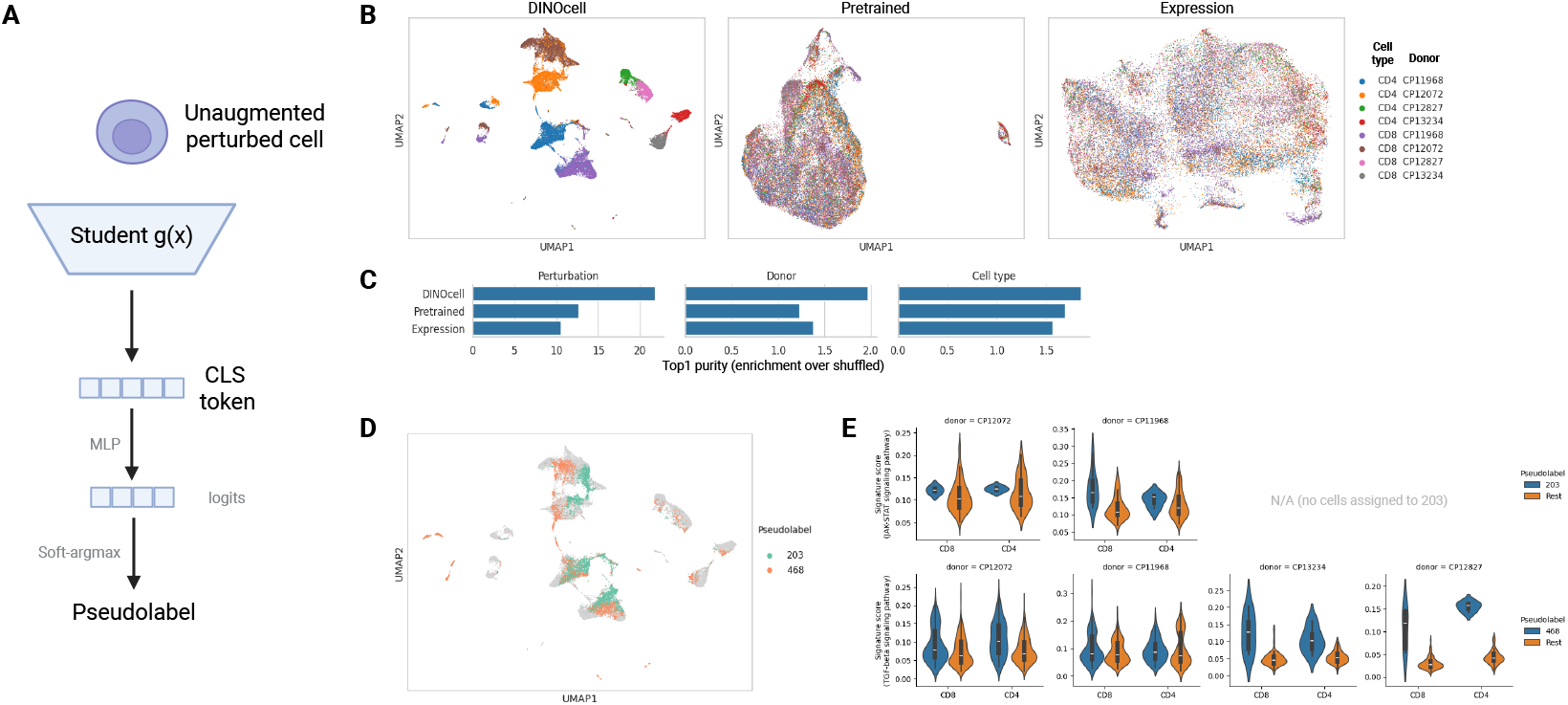
DINOcell generates semantically meaningful embeddings. A) Schematic of DINOcell embeddings: We generate embeddings by prompting the student network with unaugmented perturbed cells. The CLS embeddings represent global cell state and are ultimately used for expression regression. We examine the CLS tokens in B and C. During training, logits are computed from the CLS token and are used for aligning the latent representations between the student and teacher. This roughly corresponds to a soft-classification task for emergent clusters, termed pseudolabels. We examine pseudolabels in D and E. Note that pseudolabels are for diagnostic purposes only and are not part of the computation graph during training. B,C) DINOcell embeddings are separable by covariate groups and perturbation labels: Single cells were uniformly sampled from the gxe10 and gxe63 datasets. B: UMAP projections are shown for DINOcell embeddings (left), the base pretrained model (middle), and log-normalized expression (right), with cells colored by covariate group. DINOcell embeddings display clearer separation across covariates. C: Separability by perturbation or covariate was quantified using Top-1 purity; a k-NN classifier was trained on each representation, and the fraction of cells whose nearest neighbor shared the same label was measured relative to a shuffled baseline (n=100). D,E) DINOcell pseudolabels correspond to biologically meaningful clusters: D: Cells assigned to pseudolabels 203 and 468 are visualized on the DINOcell CLS token UMAP in B. E: Signature scores, by covariate group, for pathways corresponding to the highlighted pseudolabels. Scores are computed as the mean logFC magnitude across genes within a pathway. No cells in donors CP13234 and CP12827 were assigned to pseudolabel 203. Signature scores are higher in cells assigned to the highlighted pseudolabels compared to background.

We next evaluated whether DINOcell embeddings shared a similar structure between different T cell datasets. We reason that, within primary human T cells that differ in genetic background (e.g. donor), lineage (e.g. CD4 or CD8), or specific cell state (e.g. effector memory, central memory), a perturbation should induce broadly similar transcriptional changes in analogous pathways. Using the embeddings generated for Figure 4B, we tested whether DINOcell’s embeddings for a given perturbation in one T cell dataset were more similar to the same perturbation in another T cell dataset compared with the pretrained model. We focused on the ~ 200 perturbations that were common to both the gxe10 and gxe63 datasets. We again measured this similarity by k-NN top-1 evaluation. We found that DINOcell’s embeddings had a 28.4% increase in enrichment compared to an expression baseline and a 12.7% increase compared to masked language model pretrained embeddings (Supplementary Figure 3C). Together, these results demonstrate that DINOcell learns to separate known experimental and biological covariates (donor, lineage, cell state), while also maintaining internal structure within each covariate, reflective of the perturbation effects.

Lastly, we examined how the projection head shapes DINOcell’s representation space. This head maps the CLS embedding to a K-dimensional logit vector, which can be interpreted as pseudolabel predictions (MLP in Figure 4A; i.e., a soft classification over emergent clusters in the latent space). Under this view, cells assigned to the same pseudolabel should share semantic/biological signals and lie nearby in embedding space. If our model had experienced mode collapse, pseudolabels would correspond to the largest sources of variation within the dataset such as covariate groups.

To assess whether these pseudolabels reflect meaningful organization, we extracted the prototype logits (Figure 4A) for cells in gxe63 and asked whether their assignments captured information about the original perturbation. A UMAP of pseudobulked expression colored by pseudolabels revealed that many pseudolabels contained cells corresponding to the same perturbation across multiple covariate groups (Figure 4D). We note that, due to group centering, we would not necessarily expect pseudolabels to overlap between covariate groups. Yet, 39% (289/736) of perturbations were assigned to the same pseudolabel regardless of covariate group, suggesting that the projection head is utilizing some shared structure in the CLS token space. These shared pseudolabels frequently corresponded to perturbations targeting related pathways. For instance, pseudolabel 468 was enriched (enrichr [11], adjusted p-value = 1.8e-4) for perturbations targeting genes in the JAK-STAT pathway driving Th17 differentiation (*IL12RB1, JAK1, JAK2, JAK3, BATF, IRF4*) [10, 12, 13]. Pseudolabel 203 is enriched for genes that are functionally related to TGF*β* signaling (e.g. *TGFBR1, TGFBR2, PTEN, JUNB*; enrichr adjusted p-value = 1.4e-6) [14]. Perturbed cells corresponding to these pseudolabels also are functionally distinct from the background, as assessed by signature scores (Figure 4E). Together, these results indicate that the prototype projection head encourages DINOcell to align semantically related perturbations across contexts, grouping them into coherent latent categories rather than modeling each covariate independently.

## 3 Conclusions

Predicting cellular responses to perturbations has the potential to enable target discovery at an unprecedented scale. However, current models fail to generalize across contexts or outperform simple baselines [1, 2]. To overcome these challenges we introduced DINOcell, a weakly supervised framework that can learn robust representations of cells enabling generalization to unseen experiments.

In our results we show DINOcell can successfully predict the effects of unseen single-gene perturbations in various immortalized human cell lines and in Th17 T cells, significantly outperforming baselines. Furthermore, we demonstrate that the model learns epistatic genetic interactions by accurately predicting non-additive effects of unseen pairs of CRISPRi/KO guides. Finally, we show that representations learned by the model capture important biological variation, are robust to technical covariates, and show improved consistency across experiments.

We believe DINOcell addresses two key challenges that have limited the quality of embeddings in virtual cell models. First, we hypothesize that embeddings produced by models trained with masked token objectives may overrepresent superficial co-expression structure rather than functional relationships. Because gene expression features are highly correlated, such models can find shortcuts to reconstruct masked genes from redundant local signals. This issue has been documented in the vision domain in which image patches are often highly correlated [15]. DINOcell mitigates this by learning embeddings through a bottlenecked, multi-objective setup. Each cell is encoded into a single CLS embedding that must both align with a momentum teacher and support expression prediction through a shallow FiLM-conditioned decoder. This constrained architecture, combined with aggressive masking (as low as 256 genes in the student view), may help discourage shortcut learning and encourage more semantically meaningful embeddings.

The second challenge is that single-cell variation occurs across multiple scales, from broad differences in cell type, donor, and experimental conditions to subtle, perturbation-driven shifts in cell state. The inherently low signal-to-noise nature of scRNA-seq further obscures this fine-grained variation. We hypothesize that many models primarily capture the coarse structure of the data but struggle to represent these subtle functional effects. DINOcell addresses this challenge through two complementary strategies. First, we augment the input data by repeatedly sampling each cell and injecting Gaussian noise into expression measurements, encouraging robustness to measurement noise. Second, we apply conditional centering, which explicitly guides the model to learn representations that are invariant across covariate groups while retaining biologically relevant differences. In future work, it may be valuable to explore architectures in which such grouping is learned dynamically rather than predefined.

Together, these design choices yield functionally meaningful embeddings that share similarity between cell contexts, properties essential for building predictive and generalizable virtual cell models. Future work can focus on extending this framework to non-genetic perturbations, such as small or large molecule therapeutics and integrating multi-omic readouts. Additionally, we believe that, with some modification, our approach may be useful during pretraining as well. Ultimately, continued improvement of cell models will provide a new generalizable toolkit for discovering therapeutic targets.

## 4 Methods

### Th17 CRISPRi/KO screening workflow description (Pooled genome-wide and arrayed)

For *in vitro* Th17/Tc17 differentiation, bulk human CD4+/CD8+ T cells from 2–4 donors were activated with anti-CD3 and anti-CD28 and cultured in the presence of IL-1*β*, IL-6, IL-23 and TGF-*β* for 8–15 days. Polarization was confirmed by flow cytometry staining for the master Th17-driving transcription factor ROR*γ*t and measurement of IL-17A/F secretion. For pooled perturbations, cells were lentivirally transduced to express dCas9-KRAB and a genome-wide human CRISPRi sgRNA library, followed by antibiotic selection. For arrayed perturbations, cells were electroporated with Cas9 and sgRNAs for gene knockout. On days 7–14, cells were restimulated with Immunocult (anti-CD2/anti-CD3/anti-CD28) for 24 hours prior to flow sorting for viability and 10x scRNA-seq profiling.

### Th17 Gene selection for arrayed datasets (gxe46 / gxe63)

For gxe46, a total of 183 perturbations were queried with the following breakdown of different functional genetic edits: LOF (n=84), LOF/LOF (n=10), LOF/*IL23R* LOF (n=89). This arrayed screen targeted 89 genes including curated Th17 polarization regulators known from literature (n=26), autoimmunity clinical targets (n=35) and Th17 genome-wide CRISPRi screen hits (n=28).

For gxe63, nominations for arrayed gene perturbations were derived from analysis of a Th17 genome-wide CRISPRi screen, including genes that increased or decreased a published human Th17 gene signature score [16] and/or a general transcriptional shift score (e-dist). Essential genes were removed from the gene list anticipating little to no survival after their perturbation by CRISPR knock-out. The final perturbation list included 200 perturbations targeting a single gene and 528 targeting a combination of two different genes.

---

**Supplementary Figure 1:**
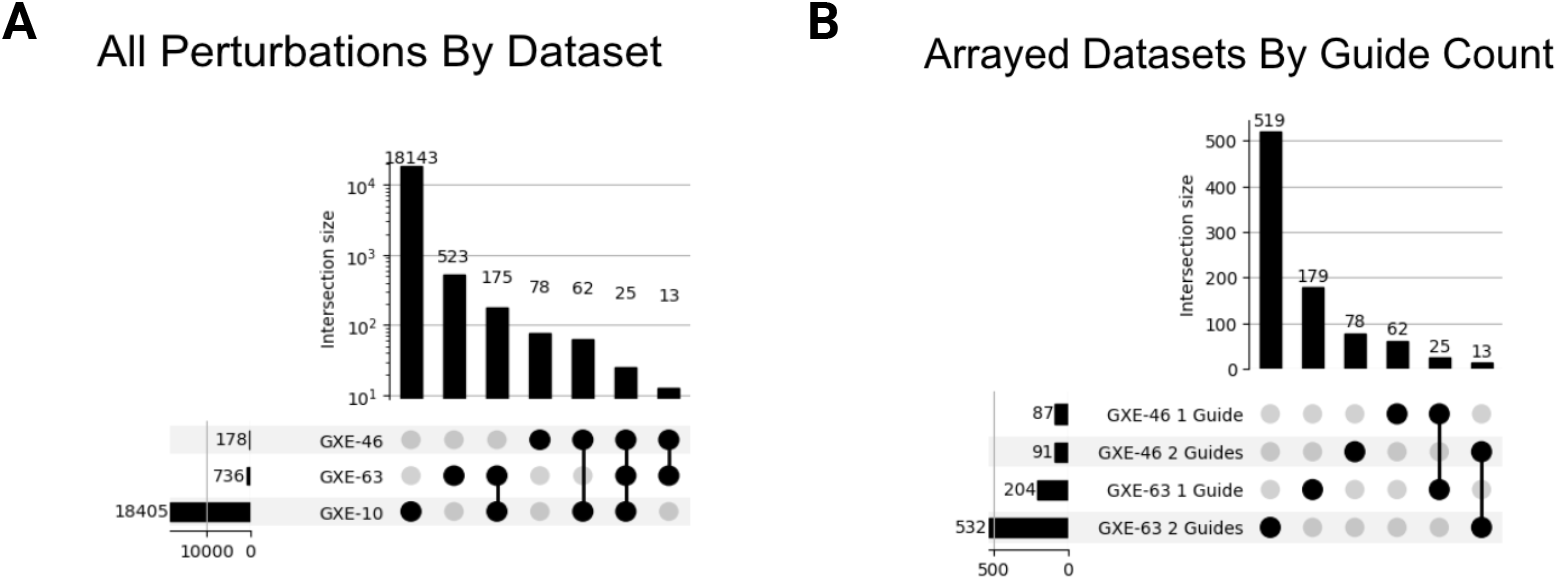
Dataset Overview. A,B) Upset plots describing number of perturbations by dataset. All datasets are CRISPRi engineered Th17 polarized T-cells. gxe10 is a genome-wide perturb-seq dataset. gxe46 and gxe63 were generated in a plate-based assay. C)Describes the overlap of CRISPRi perturbations across all three datasets. D)Describes the overlap of CRISPRi perturbations for the plate-based assays for perturbations targeting either a single gene or a combination of two different genes. gxe10, the genome-wide dataset, only includes single guide perturbations.

**Supplementary Figure 2:**
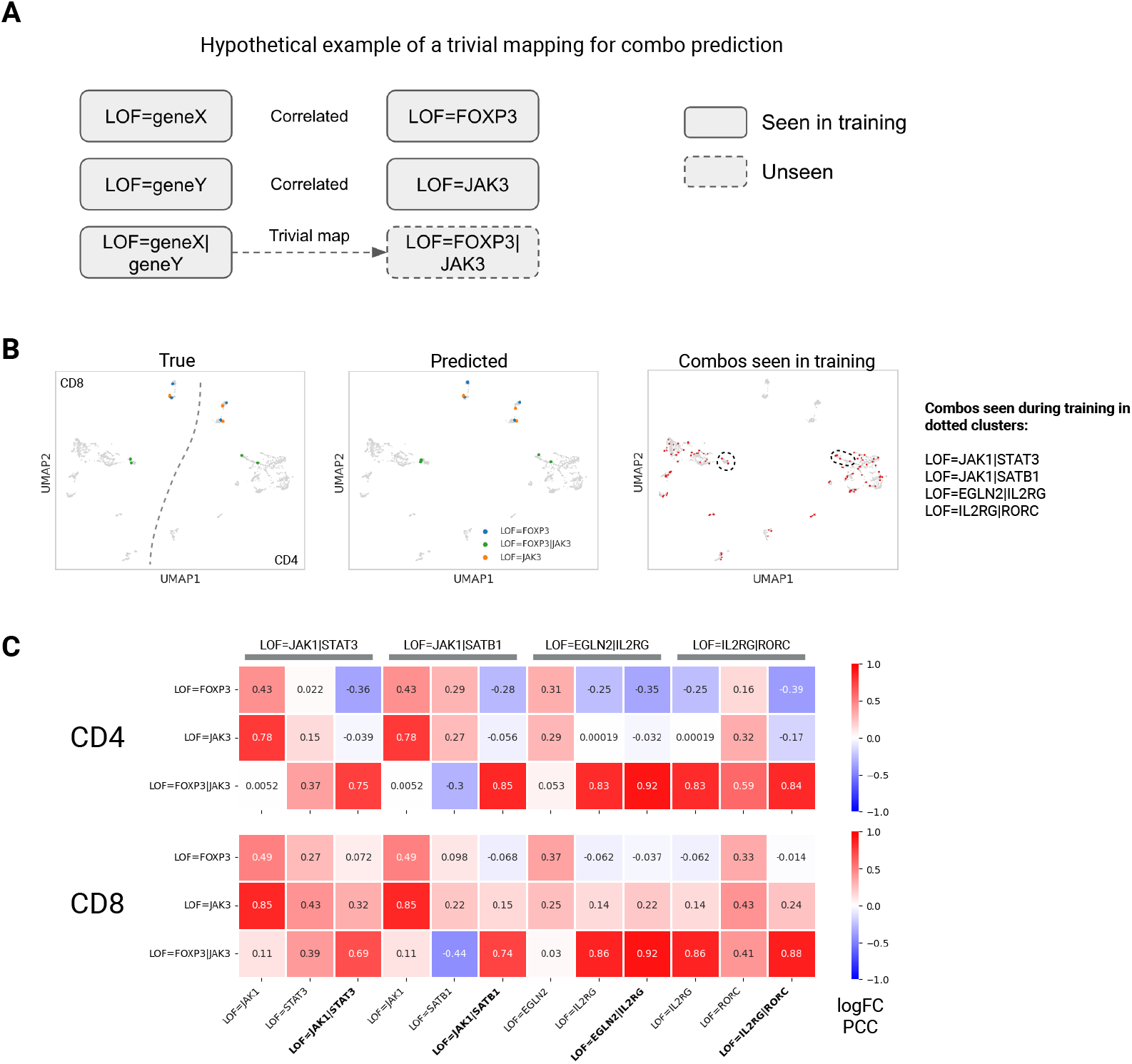
A hypothetical example showcasing a trivial mapping for combination prediction. A)For some perturbations, models may learn to predict combinations trivially. For example, if LOF=*FOXP3*|*JAK3* was seen during training and its component perturbations are correlated with those of another combination, the model may effectively memorize these associations, resembling a one-hot encoding. B) Identification of 4 training combinations that cluster with LOF=*FOXP3*|*JAK3*: UMAP projections of pseudobulked gxe63 expression (by perturbation and covariate group) are shown in gray. (Left) LOF=*FOXP3*|*JAK3* and its single-gene components are highlighted for both CD4 and CD8 cells (separated by dotted line). (Middle) DINOcell-predicted expression shown in the same space. (Right) 4 combinations were found to share the same clusters (dotted lines) with LOF=*FOXP3*|*JAK3* in either CD4 or CD8 cells. C) None of the 4 training combinations offer a trivial mapping for predicting LOF=*FOXP3*|*JAK3*: Each of the 4 combinations in B were assessed to determine whether they fit the hypothetical schematic in A. Each square displays the logFC PCC between: LOF=*FOXP3*|*JAK3* and its singles, and each of the 4 combinations and their singles.

**Supplementary Figure 3:**
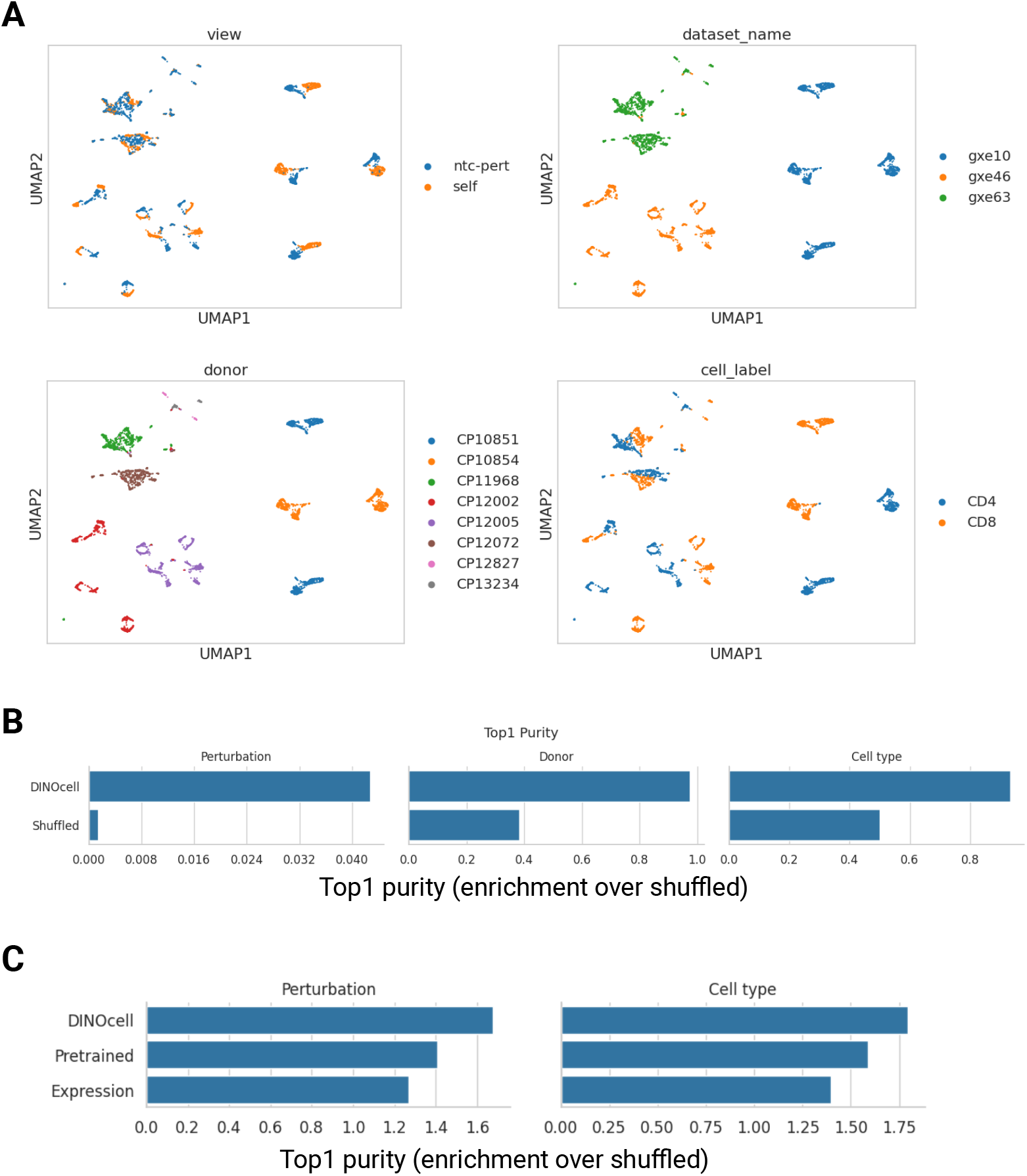
CLS embeddings align between conditional and observed views and across datasets. A) Single cells were sampled from the GXE datasets and used to generate paired embeddings representing either conditional and observed views. The resulting embeddings were plotted on a UMAP (left). Both views produced embeddings that clustered with the same covariate groups. B) Separability by perturbation or covariate was quantified using Top-1 purity; a k-NN classifier was trained on each representation, and the fraction of cells whose nearest neighbor shared the same label was measured relative to a shuffled baseline (n=100). C) CLS embeddings align across datasets: 20 cells are sampled per covariate group for each of the 200 perturbations in common between gxe10 and gxe63. We compute top-1 purity scores, normalized by a shuffled baseline, between datasets. More specifically, for all cells in one dataset, we assess the frequency that the closest neighbor in the other dataset is the same perturbation or cell type.

## Footnotes

1 Control cells in Replogle-Nadig are untreated cells. In GXE cohorts these were treated with non-targeting CRISPR guide RNA (NTC).

## References

[1] Constantin Ahlmann-Eltze, Wolfgang Huber, and Simon Anders. Deep-learning-based gene perturbation effect prediction does not yet outperform simple linear baselines. Nature Methods, 22(8):1657–1661, 2025.

[2] Daniel R. Wong, Abby S. Hill, and Rob Moccia. Simple controls exceed best deep learning algorithms and reveal foundation model effectiveness for predicting genetic perturbations. Bioinformatics, 41(6):btaf317, 2025.

[3] Mathilde Caron, Hugo Touvron, Ishan Misra, Hervé Jégou, Julien Mairal, Piotr Bojanowski, and Armand Joulin. Emerging properties in self-supervised vision transformers. arXiv preprint 2104.14294, 2021.

[4] Haotian Cui, Chloe Wang, Hassaan Maan, Kuan Pang, Fengning Luo, Nan Duan, and Bo Wang. scGPT: toward building a foundation model for single-cell multi-omics using generative ai. Nature Methods, 21(8):1470–1480, 2024.

[5] Ethan Perez, Florian Strub, Harm de Vries, Vincent Dumoulin, and Aaron Courville. FiLM: visual reasoning with a general conditioning layer. arXiv preprint 1709.07871, 2017.

[6] Yiqun Chen and James Zou. GenePT: A simple but effective foundation model for genes and cells built from chatgpt. bioRxiv, 2024. Preprint.

[7] Dan Hendrycks and Kevin Gimpel. Gaussian error linear units (GELUs). arXiv preprint 1606.08415, 2016.

[8] Joseph M. Replogle, Reuben A. Saunders, Angela N. Pogson, Jeffrey A. Hussmann, Alexander Lenail, Alina Guna, Lauren Mascibroda, Eric J. Wagner, Karen Adelman, Gila Lithwick-Yanai, Nika Iremadze, Florian Oberstrass, Doron Lipson, Jessica L. Bonnar, Marco Jost, Thomas M. Norman, and Jonathan S. Weissman. Mapping information-rich genotype-phenotype landscapes with genome-scale Perturb-seq. Cell, 185(14):2559–2575.e28, 2022.

[9] Ajay Nadig, Joseph M. Replogle, Angela N. Pogson, Mukundh Murthy, Steven A. McCarroll, Jonathan S. Weissman, Elise B. Robinson, and Luke J. O’Connor. Transcriptome-wide analysis of differential expression in perturbation atlases. Nature Genetics, 57(5):1228–1237, 2025.

[10] Peng Li, Rosanne Spolski, Wei Liao, Lu Wang, Theresa L. Murphy, Kenneth M. Murphy, and Warren J. Leonard. BATF–JUN is critical for IRF4-mediated transcription in t cells. Nature, 490(7421):543–546, 2012.

[11] Maxim V. Kuleshov, Matthew R. Jones, Andrew D. Rouillard, Nicolas F. Fernandez, Qiaonan Duan, Zichen Wang, Simon Koplev, Sherry L. Jenkins, Kathleen M. Jagodnik, Alexander Lachmann, Michael G. McDermott, Caroline D. Monteiro, Gregory W. Gundersen, and Avi Ma’ayan. Enrichr: a comprehensive gene set enrichment analysis web server 2016 update. Nucleic Acids Research, 44(W1):W90–W97, 2016.

[12] Barbara U. Schraml, Kai Hildner, Wataru Ise, Wan-Ling Lee, Whitney A.-E. Smith, Ben Solomon, Gurmukh Sahota, Julia Sim, Ryuta Mukasa, Saso Cemerski, Robin D. Hatton, Gary D. Stormo, Casey T. Weaver, John H. Russell, Theresa L. Murphy, and Kenneth M. Murphy. The AP-1 transcription factor batf controls Th17 differentiation. Nature, 460(7253):405–409, 2009.

[13] Theresa L. Murphy, Roxane Tussiwand, and Kenneth M. Murphy. Specificity through cooperation: BATF–IRF interactions control immune-regulatory networks. Nature Reviews Immunology, 13(7):499–509, 2013.

[14] Joan Massagué and Dean Sheppard. TGF-β signaling in health and disease. Cell, 186(19):4007–4037, 2023.

[15] Kaiming He, Xinlei Chen, Saining Xie, Yanghao Li, Piotr Dollár, and Ross Girshick. Masked autoencoders are scalable vision learners. arXiv preprint 2111.06377, 2021.

[16] Eddie Cano-Gamez, Blagoje Soskic, Theodoros I. Roumeliotis, Ernest So, Deborah J. Smyth, Marta Baldrighi, David Willé, Nikolina Nakic, Jorge Esparza-Gordillo, Christopher G. C. Larminie, Paola G. Bronson, David F. Tough, Wendy C. Rowan, Jyoti S. Choudhary, and Gosia Trynka. Single-cell transcriptomics identifies an effectorness gradient shaping the response of cd4+ t cells to cytokines. Nature Communications, 11(1):1801, 2020.

